# Distinct nigral and brainstem pathology markers map onto separable subthalamic electrophysiological signatures in Parkinson’s disease

**DOI:** 10.64898/2026.07.17.739149

**Authors:** A Delgado-Sanchez, L Andrews, P Hayton, C Craig, A Macerollo, J Cortes-Gutierrez, S Martin, R Somervail, A Azimi, MLTM Müller, L Parkes, H Haroon, M Bergamino, SA Kotz, M Silverdale, NJ Trujillo-Barreto, NJ Ray

**Affiliations:** Manchester Metropolitan University, School of Psychology, Translational and Computational Neurosciences, Manchester, UK; The Walton Centre NHS Foundation Trust for Neurology and Neurosurgery, Liverpool, UK; Institute of Systems, Molecular and Integrative Biology, The University of Liverpool, UK; Department of Neurology, Manchester Centre for Clinical Neurosciences, Manchester Academic Health Science Centre, University of Manchester, Manchester, United Kingdom; Division of Neuroscience & Experimental Psychology, School of Biological Sciences, Faculty of Biology, Medicine and Health, University of Manchester, UK; Barrow Neurological Institute, Phoenix AZ, USA; Maastricht University, Faculty of Psychology and Neuroscience, Universiteitssingel 40, 6200 MD, Maastricht, The Netherlands; Max Planck Institute for Human Cognitive and Brain Sciences, Department of Neurology, Stephanstrasse 1A, 04103 Leipzig, Germany

**Author notes:** shared senior authorship (M Silverdale, NJ Trujillo-Barreto, NJ Ray).

## Abstract

Subthalamic local field potentials (LFPs) are increasingly used as physiomarkers of the symptomatic state in Parkinson’s disease, but their relationship to the underlying neurodegenerative pathology remains unclear. Here, we combined OFF-medication subthalamic LFP recordings with quantitative MRI markers of nigral and brainstem pathology in 33 people with Parkinson’s disease. Distinct pathological markers mapped onto dissociable electrophysiological components. Substantia nigra pars compacta susceptibility was associated with increased occupancy, duration and rate of low-β bursts, whereas nigral free water was associated with greater low-frequency aperiodic offset and a steeper slope. Pedunculopontine nucleus free-water-corrected axial diffusivity was selectively associated with high-frequency aperiodic activity, and this relationship strengthened with increasing nigral susceptibility, consistent with dopaminergic-state-dependent influences of extranigral pathology on subthalamic physiology. Only low-frequency aperiodic offset was also associated with contralateral bradykinesia. These findings indicate that the subthalamic LFP is not a unitary readout of dopamine loss or motor state, but an integrated physiological signal in which pathology across interconnected systems is expressed through separable oscillatory and aperiodic components. Chronically implanted devices may therefore provide physiological readouts of underlying disease biology alongside control signals for adaptive therapy.

## 3. Introduction

Subthalamic local field potentials can now be recorded chronically and acted upon therapeutically, yet what these signals reveal about the underlying biology of Parkinson’s disease remains poorly understood. Recordings from deep brain stimulation (DBS) leads in the subthalamic nucleus (STN), a key basal ganglia node and major therapeutic target, have established exaggerated β-band synchronisation (13–30 Hz) as a prominent electrophysiological feature of the parkinsonian state. STN β power and β-burst activity are elevated following medication withdrawal, relate most consistently to bradykinesia and rigidity, and are reduced by levodopa and effective STN stimulation ^1–10^. These properties have positioned β activity as a marker of the symptomatic parkinsonian state and a control signal for adaptive DBS, but its relationship to the neurodegenerative processes that generate circuit dysfunction remains unclear. Showing that specific LFP features reflect underlying neurodegeneration in humans would extend chronically implanted DBS systems beyond symptom-responsive therapy, enabling them to monitor disease progression and provide objective physiological endpoints for trials of neuroprotective and disease-modifying interventions.

A key unresolved question is therefore how pathological STN activity develops in relation to progressive neurodegeneration. Because human LFP recordings are usually obtained only after patients have reached the advanced disease stage required for DBS, they cannot readily determine whether β abnormalities emerge early, increase with accumulating pathology, or predominantly reflect the symptomatic state at implantation. Longitudinal studies in dopamine-depleted and α-synuclein overexpression models show that increases in STN β power and prolonged β bursting can emerge alongside nigrostriatal pathology and track dopaminergic cell loss, reduced striatal fibre density and motor decline ^11–13^. However, few human datasets combine in vivo markers of Parkinson’s disease pathology with STN LFP recordings, leaving it unclear which pathological profiles are expressed in the electrophysiological features. Hofman et al.^13^ found a link between β activity, striatal dopamine uptake and motor impairment in a small sample of people with Parkinson’s disease, but the study was not powered to test direct relationships between dopaminergic pathology and STN electrophysiology. Advances in MRI markers of regional neurodegeneration now provide an opportunity to address this gap by testing how distinct patterns of pathology are expressed in human basal ganglia physiology, thereby refining the biological interpretation of STN LFP features.

Importantly however, narrow-band β oscillations represent only one component of the STN LFP signal. Neural power spectra also contain a broadband, non-oscillatory component whose power decreases with increasing frequency and which has historically been treated as background activity. This aperiodic structure is increasingly recognised as physiologically informative and can be parameterised by its offset and slope or exponent ^14–18^. The offset broadly reflects the overall magnitude of broadband population activity, whereas the slope describes how rapidly power decays across frequencies and has been related to underlying synaptic and population dynamics. Importantly, the physiological interpretation of aperiodic parameters may also depend on the frequency range over which they are estimated. For example, in the parkinsonian STN, exponents estimated over 40–90 Hz in humans have been linked to neuronal firing and excitation–inhibition balance. Although their precise biophysical interpretation remains debated, these parameters provide information about circuit activity that is not captured by narrow-band oscillatory power alone ^15,17–19^. This may be especially relevant in Parkinson’s disease, where basal ganglia dysfunction involves altered firing rates and spike timing as well as exaggerated oscillatory synchronisation ^20–22^. Emerging evidence indicates that STN aperiodic activity varies with motor impairment, clinical asymmetry and dopaminergic lesion severity, and may therefore capture dimensions of parkinsonian circuit dysfunction not indexed by β activity ^23,24^.

Quantitative MRI provides a means of testing whether distinct STN electrophysiological features are associated with different in vivo markers of nigral pathology. Free water within the substantia nigra is elevated in Parkinson’s disease and is commonly interpreted as reflecting microstructural tissue disruption, extracellular expansion and neurodegenerative change ^25–27^. Iron-sensitive susceptibility measures provide an in vivo index of abnormal iron accumulation in the substantia nigra ^28,29^, while post-mortem and experimental evidence suggests that iron accumulation may be both a consequence of tissue degeneration and a contributor to oxidative and cellular injury ^30,31^. Diffusion tractography provides a complementary pathway-level measure of nigrostriatal organisation, offering an in vivo human counterpart to the striatal fibre-density measures linked to β abnormalities in longitudinal animal models ^13^. Together, these regional and tract-based measures offer a way to test how variation in nigral tissue properties and nigrostriatal pathway integrity is expressed in downstream human basal ganglia activity.

Yet nigral pathology is unlikely to account fully for the heterogeneity of STN activity, because Parkinson’s disease also affects other systems capable of shaping basal ganglia physiology. One such system is the pedunculopontine nucleus (PPN), a brainstem structure implicated in gait, arousal and motor control. Through widespread reciprocal connections with basal ganglia circuits, the PPN is well positioned to influence STN physiology ^32–35^. Human intracranial recordings further support functional communication between the PPN and STN, with similar locally generated evoked responses observed in both structures, consistent with transmission of physiologically meaningful activity through PPN–STN connectivity ^36^. Importantly, preclinical evidence suggests that the nature of this influence depends on dopaminergic integrity. PPN lesions induce STN hyperactivity in dopamine-intact rodents but reduce elevated STN firing after dopamine depletion ^37,38^. In addition, low-frequency PPN-region stimulation selectively reduces elevated STN firing and β activity in 6-OHDA-lesioned, but not sham-lesioned rodents ^38.5^. These opposing effects indicate that PPN pathology may alter STN physiology differently depending on the integrity of the nigrostriatal system, creating an interaction between dopaminergic and extranigral degeneration in the emergence of pathological basal ganglia activity.

This hypothesis can now be examined in vivo using diffusion MRI measures sensitive to PPN tissue properties. PPN free water and free-water-corrected axial diffusivity (cAD) have been associated with disease-related changes in Parkinson’s disease ^39,40^. Considered alongside nigral free water and susceptibility, these measures allow us to test what distinct STN LFP features reveal about the underlying biology of Parkinson’s disease: whether they provide specific readouts of nigral and extranigral pathology, and whether their expression is shaped by interactions between these degenerating systems.

Here, we combined OFF-medication STN LFP recordings with quantitative MRI measures of ipsilateral nigral and PPN tissue properties and nigrostriatal pathway integrity. We tested whether β power, β-burst dynamics and aperiodic spectral features were differentially associated with these imaging markers, whether relationships between PPN microstructure and STN physiology depended on nigral pathology, and whether electrophysiological and imaging measures related to contralateral motor severity. We hypothesised that STN LFP features would be associated with markers of nigral and extranigral pathology, and (given the preclinical literature discussed above) that the electrophysiological expression of PPN tissue change would be most evident in those with more significant nigral pathology.

## 4. Results

### 4.1 Overview of STN-LFP and multimodal MRI analysis

For an overview of the study see Figure 1. To test whether STN electrophysiological features are associated with in vivo markers of nigral and brainstem pathology, we combined OFF-medication STN local field potential recordings with pre-operative quantitative MRI measures of SNc and PPN tissue properties in 33 people (from 45 recruited individuals) with Parkinson’s undergoing DBS. Electrophysiological outcomes comprised β-band power, β-burst dynamics and broadband aperiodic features. Since the physiological interpretation of aperiodic parameters may also depend on the frequency range over which they are estimated, we examined aperiodic structure separately over lower (4–39 Hz) and higher (40–80 Hz) frequency ranges.

**Figure 1.**
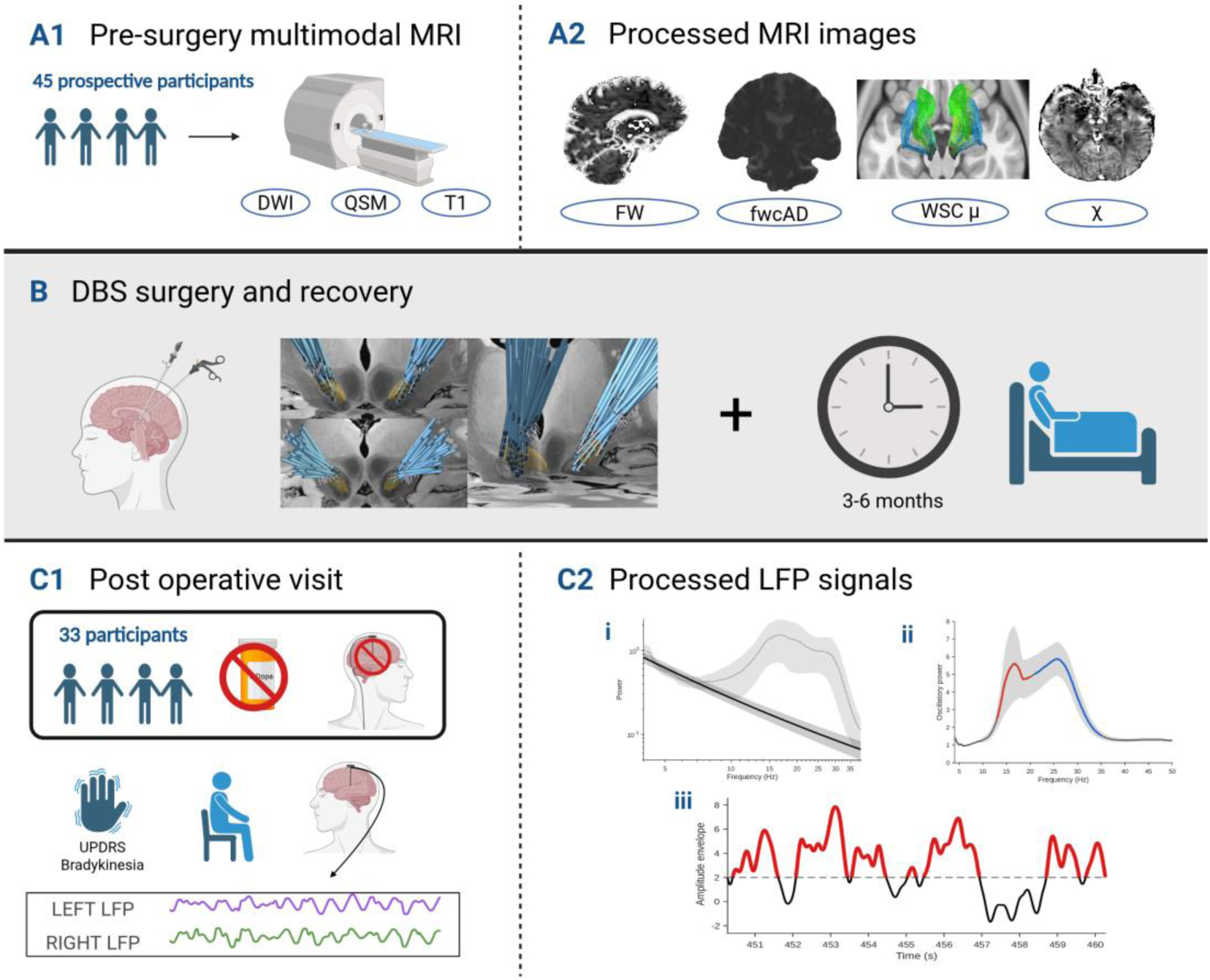
Study overview and derivation of MRI and electrophysiological measures. **(A1)** Forty-five prospective participants underwent multimodal MRI before deep brain stimulation (DBS) surgery. **(A2)** Representative pre-processed MRI maps included free water (free water), free water-corrected axial diffusivity (cAD), weighted streamline density (μ), and magnetic susceptibility (χ) maps; **(B)** Following MRI acquisition, participants underwent DBS implantation as part of routine clinical care (greyed panel). Participants recovered for approximately 3–6 months, depending on clinical need. Image shows the location of implanted DBS electrodes overlaid on a template brain; **(C1)** Thirty-three participants completed the post-operative research visit after overnight withdrawal of levodopa medication and with DBS switched OFF. Clinical motor assessments and resting-state subthalamic nucleus (STN) local field potential (LFP) recordings were obtained; **(C2)** Resting-state LFP recordings were used to derive three classes of electrophysiological measures. **(i)** *Aperiodic activity*: plot shows the grand-average power spectrum (grey dotted line) and the fitted aperiodic component (black solid line); shaded regions indicate the standard error of the mean (SEM). Aperiodic parameters (offset and slope) were extracted separately for the 4–39 Hz and 40–80 Hz frequency ranges; the figure displays the 4–39 Hz fit. **(ii)** *Oscillatory activity*: the grand-average oscillatory spectrum following removal of the aperiodic component. The shaded region represents the SEM. Low-β (red) and high-β (blue) frequency bands were analysed separately. **(iii)** *Burst analysis*: representative burst detection from a single participant. Bursts were defined as periods during which the robust z-scored amplitude envelope exceeded a threshold of 2. From the detected bursts, burst rate, occupancy, median duration, amplitude, and inter-burst interval were calculated.

Hemisphere-level associations were tested using mixed-effects models that accounted for repeated observations (i.e. left and right hemisphere) within participants and adjusted for age, residual OFF-medication levodopa exposure and the relevant whole-brain grey-matter MRI measure. False discovery rate (FDR) correction was applied within prespecified MRI-electrophysiology analysis families. All principal associations were subsequently evaluated using prespecified sensitivity analyses addressing data quality, influential observations and model specification.

### 4.2 Nigral free water and susceptibility map onto distinct STN electrophysiological signatures

We first tested whether MRI markers of nigral tissue pathology and nigrostriatal tract integrity were associated with STN electrophysiology. SNc free water, SNc susceptibility and nigrostriatal tract measures were examined in separate models. Free-water and susceptibility analyses were additionally adjusted for the corresponding whole-brain grey-matter measure. Descriptive statistics and model-specific sample sizes (following quality control exclusions, described in Methods) are reported in Supp Table 1.

SNc susceptibility was selectively associated with low-β burst dynamics rather than aperiodic spectral features. It was not associated with aperiodic offset or slope in either the 4–39 Hz or 40–80 Hz ranges. Instead, higher SNc susceptibility was selectively associated with low-β burst dynamics, predicting greater low-β percentage burst time (β = 0.59, t = 5.15, p = 5.87 × 10⁻⁶, q = 4.69 × 10⁻⁵), longer low-β median burst duration (β = 0.51, t = 3.69, p = .0006, q = .0024), and higher low-β burst rate (β = 0.33, t = 2.57, p = .014, q = .036). No corrected associations were observed for high-β burst metrics, β-band oscillatory spectral mass, or aperiodic measures.

In contrast, higher SNc free water was associated with alterations in low-frequency aperiodic activity in the ipsilateral STN. Specifically, SNc free water was associated with a steeper 4–39 Hz aperiodic slope (β = 0.36, t = 2.68, p = .001, q = .038) and a higher 4–39 Hz aperiodic offset (β = 0.26, t = 2.42, p = .019, q = .038). In contrast, SNc free water was not associated with 40–80 Hz aperiodic activity or aperiodic-adjusted β-band oscillatory spectral mass. See Figure 2a-e.

**Figure 2:**
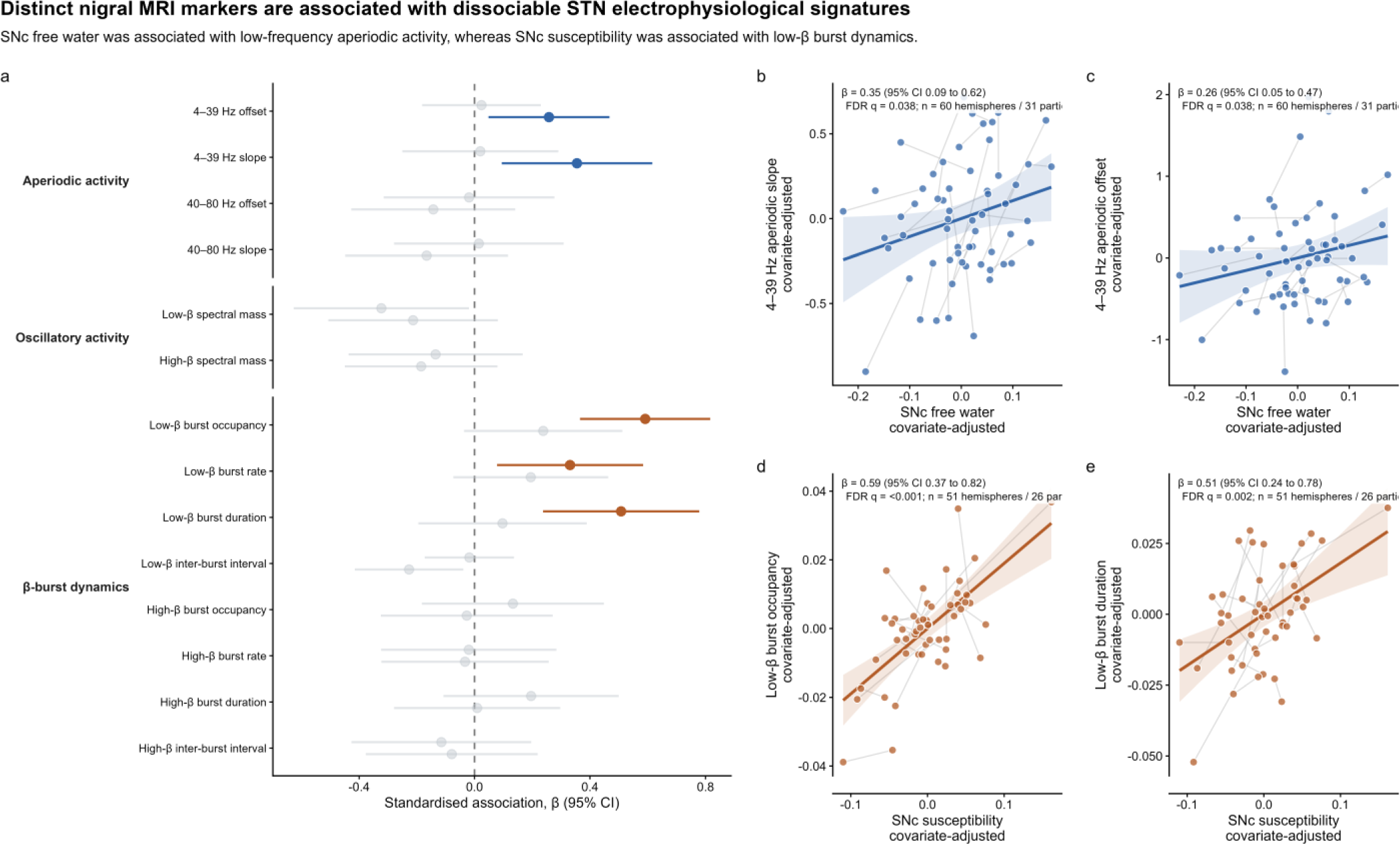
Distinct nigral MRI markers are associated with dissociable OFF-medication STN electrophysiological signatures. **a**, Standardised mixed-effects model associations between substantia nigra pars compacta (SNc) free water or susceptibility and prespecified STN electrophysiological features. Points indicate standardised regression coefficients and horizontal lines indicate 95% confidence intervals; coloured points denote associations surviving false discovery rate correction. **b,c**, Covariate-adjusted hemisphere-level data showing that higher SNc free water was associated with steeper 4–39 Hz aperiodic slope and greater 4–39 Hz aperiodic offset. **d,e**, Covariate-adjusted hemisphere-level data showing that higher SNc susceptibility was associated with greater low-β burst occupancy and longer low-β burst duration. Points represent individual hemispheres and thin grey lines connect paired hemispheres from the same participant. Coloured lines and shaded bands show ordinary least-squares fits and 95% confidence intervals.

Nigrostriatal tract metrics show no FDR-corrected associations with STN electrophysiological features

Sensitivity analyses supported the robustness of the primary findings, indicating that the observed associations were not attributed to model instability, multicollinearity or quality-control threshold selection. Importantly, matched-sample analyses restricted to participants with both free-water and susceptibility measures available confirmed the dissociation between free water and susceptibility (see Supplemental section 1)

These findings suggest that local nigral tissue markers are associated with separable electrophysiological signatures within the STN. SNc susceptibility was preferentially associated with the temporal organisation of low-β bursting, while SNc free water was preferentially associated with low-frequency aperiodic activity.

### 4.2a PPN microstructure relates to high-frequency aperiodic STN activity

We next tested whether MRI markers of PPN tissue properties were associated with STN electrophysiology. PPN cAD and free water were examined as predictors of the same LFP features described above. As above, primary models adjusted for age, estimated residual OFF-medication levodopa exposure and the corresponding whole-brain grey-matter MRI measure. To assess whether associations were independent of concurrent nigral tissue variation, we ipsilateral SNc free water and susceptibility were added as covariates in separate models.

Higher PPN cAD was selectively associated with high-frequency aperiodic STN activity, with no corresponding associations with oscillatory β power or β-burst measures. In primary whole-brain-adjusted models, higher PPN cAD was associated with greater 40–80 Hz aperiodic offset (β = 0.349, *t* = 2.87, *p* = 0.006, *q* = 0.010) and steeper 40–80 Hz aperiodic slope (β = 0.350, *t* = 2.85, *p* = 0.006, *q* = 0.010) see Figure 3a-c. Both associations remained directionally and statistically consistent after separate adjustment for ipsilateral SNc free water or susceptibility, with comparable effect sizes across models (β = 0.31–0.36, *p* = 0.008–0.018, *q* = 0.017–0.036). The findings were also retained in a modality-matched SNc cAD model and in leave-one-hemisphere-out sensitivity analyses (Supplementary Section 2).

**Figure 3.**
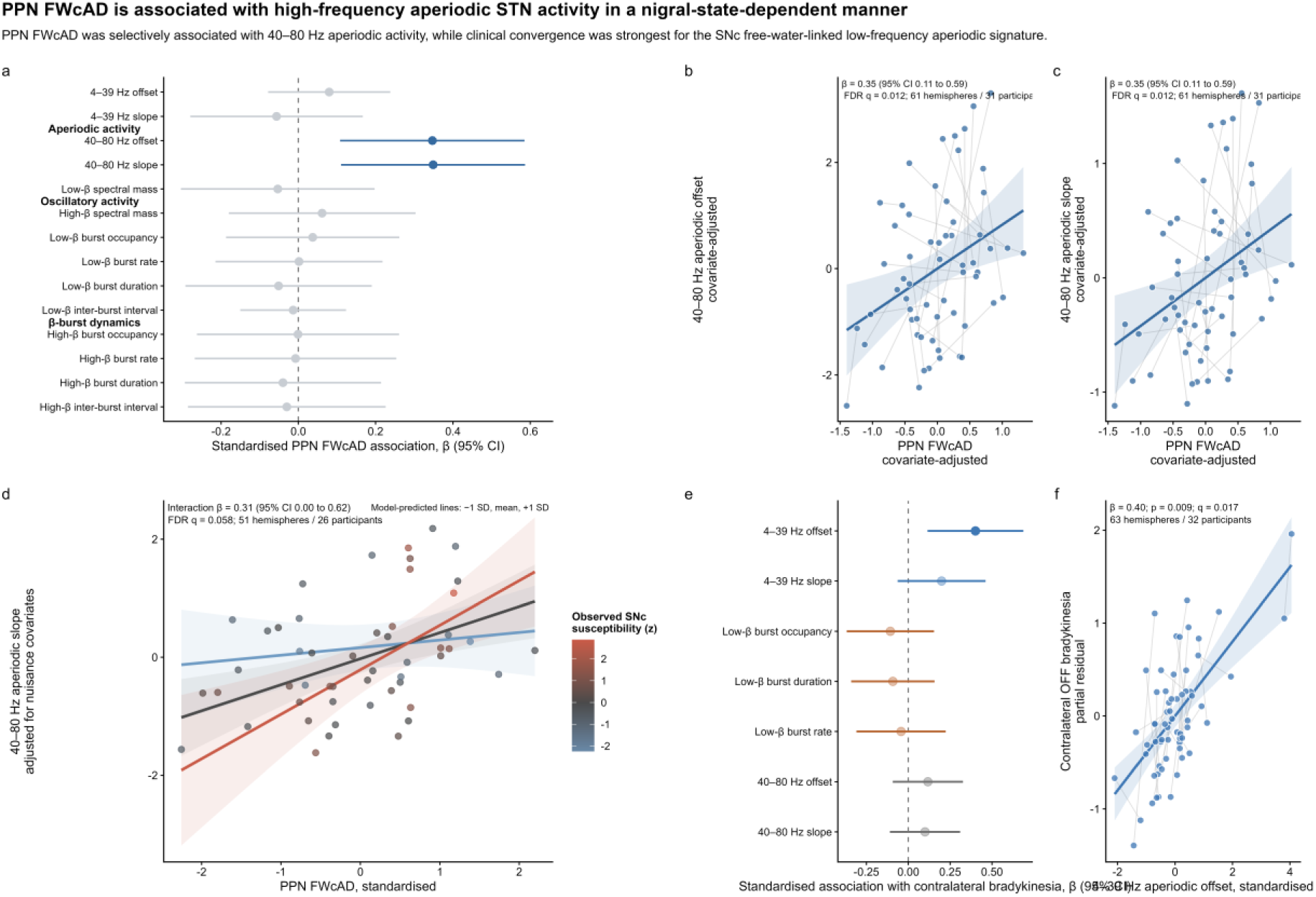
PPN microstructure is associated with high-frequency aperiodic STN activity in a nigral-state-dependent manner. a, Standardised mixed-effects model associations between pedunculopontine nucleus free-water-corrected axial diffusivity (PPN cAD) and prespecified subthalamic nucleus (STN) electrophysiological features. Points indicate standardised regression coefficients and horizontal lines indicate 95% confidence intervals. Blue points denote associations surviving false-discovery-rate correction; grey points denote non-significant associations. b,c, Covariate-adjusted hemisphere-level data showing the associations between PPN cAD and 40–80 Hz aperiodic offset (b) and slope (c). Points represent individual data points and thin grey lines connect paired hemispheres from the same participant. Blue lines and shaded bands show ordinary least-squares fits and 95% confidence intervals for visualisation. d, Interaction between PPN cAD and ipsilateral substantia nigra pars compacta (SNc) susceptibility in predicting 40–80 Hz aperiodic slope. The association between PPN cAD and aperiodic slope became stronger with increasing SNc susceptibility. The coloured line and ribbon represent model-predicted associations at observed levels of SNc susceptibility; points show covariate-adjusted hemisphere-level observations. e, Standardised mixed-effects model associations between MRI-linked STN electrophysiological features and contralateral OFF-medication bradykinesia. Points indicate standardised regression coefficients and 95% confidence intervals. f, Covariate-adjusted hemisphere-level relationship between 4–39 Hz aperiodic offset and contralateral OFF-medication bradykinesia. Points represent individual hemispheres and thin grey lines connect paired hemispheres from the same participant; the blue line and shaded band show the ordinary least-squares fit and 95% confidence interval.

PPN free water showed an inverse but less robust relationship with the same high-frequency aperiodic measures. In primary whole-brain-adjusted models, higher PPN free water was nominally associated with lower 40–80 Hz aperiodic offset and slope, but neither association survived FDR correction (β = −0.27 to −0.26, *p* = 0.038, *q* = 0.077). The associations became stronger after adjustment for SNc susceptibility, but were attenuated after adjustment for SNc free water. Thus, the evidence for an independent PPN free-water association was weaker, and sensitive to the nigral MRI covariate included.

### 4.2b Nigral susceptibility moderates the association between PPN free water-corrected AD and high-frequency aperiodic STN activity

Motivated by preclinical evidence that the effect of PPN dysfunction on STN activity depends on dopaminergic integrity, with PPN lesions reducing STN hyperactivity and β oscillatory activity after dopamine depletion ^37,38^, we tested whether ipsilateral nigral pathology moderated the association between PPN cAD and 40–80 Hz aperiodic activity. We specifically predicted that greater nigral pathology would strengthen the positive association between PPN cAD and aperiodic slope, because steeper slopes have been proposed to reflect relatively greater inhibitory influence within population activity ^24^. SNc susceptibility was selected as the primary moderator because it showed the clearest associations with STN electrophysiological features previously linked to dopaminergic dysfunction, particularly low-β burst dynamics.

The PPN cAD × SNc susceptibility interaction term was positive and significant for both 40–80 Hz aperiodic offset (β = 0.31, t = 1.97, one-tailed (consistent with the pre-specified directional hypothesis) p = .027) and 40–80 Hz aperiodic slope (β = 0.31, t = 1.95, one-tailed p = .029). Interaction plots indicated that the positive association between PPN cAD and high-frequency aperiodic activity became progressively stronger with increasing SNc susceptibility (see Figure 3d). This moderation effect was not observed for SNc free water or SNc cAD, or for PPN free water interactions, consistent with specificity for the combination of PPN cAD and SNc susceptibility.

Leave-one-hemisphere-out analyses confirmed that the interaction remained positive in all refitted models and remained significant in 98/102 refits for both outcomes (see Supplemental section 2).

### 4.3 MRI-linked STN electrophysiological signatures and contralateral bradykinesia

Motivated by recent evidence that STN aperiodic activity relates more consistently to motor impairment than β activity alone ^23^, we tested whether the MRI-linked electrophysiological signatures were also associated with contralateral OFF-medication bradykinesia. Based on previous data linking low β spectral mass with motor symptoms ^1–6,23^, and the preceding MRI-LFP analyses, we carried forward eight electrophysiological features: 4–39 Hz aperiodic offset and slope, reflecting the SNc free-water-associated low-frequency aperiodic signature; low-beta burst percentage time, median burst duration and burst rate, reflecting the SNc susceptibility-associated low-beta burst signature; 40–80 Hz aperiodic offset and slope, reflecting the PPN cAD-associated high-frequency aperiodic signature, and low β spectral mass .

In single-predictor mixed models, the clearest association with bradykinesia was observed for the SNc free-water-associated low-frequency aperiodic signature. Higher 4–39 Hz aperiodic offset in the STN was associated with worse contralateral OFF-medication bradykinesia (β = 0.401, t = 2.74, p = .007, q = .017). None of the low-beta spectral mass or burst measures, or high-frequency aperiodic measures showed significant associations with contralateral bradykinesia (Figure 2 e, f).

We additionally tested whether representative features from the three MRI-linked electrophysiological signatures, together with low-β spectral mass, contributed independent explanatory value. In an exploratory mixed-effects model simultaneously including 4–39 Hz aperiodic offset, low-β burst occupancy, low-β spectral mass and 40–80 Hz aperiodic offset, only 4–39 Hz aperiodic offset showed a nominal association with contralateral bradykinesia (β = 0.322, t = 2.41, p = .019), although this marginally missed correction across the four electrophysiological terms (q = .077). The remaining terms were not significant, and collinearity was low (maximum VIF = 1.58)., indicating that the absence of significant effects for the β-related and high-frequency aperiodic terms was not explained by redundancy between predictors.

Direct MRI–symptom models provided little evidence for simple linear relationships between structural markers and contralateral bradykinesia. None of the MRI predictors showed nominal associations with contralateral OFF-medication bradykinesia.

For a summary of all results reported above, see Figure 4.

**Figure 4.**
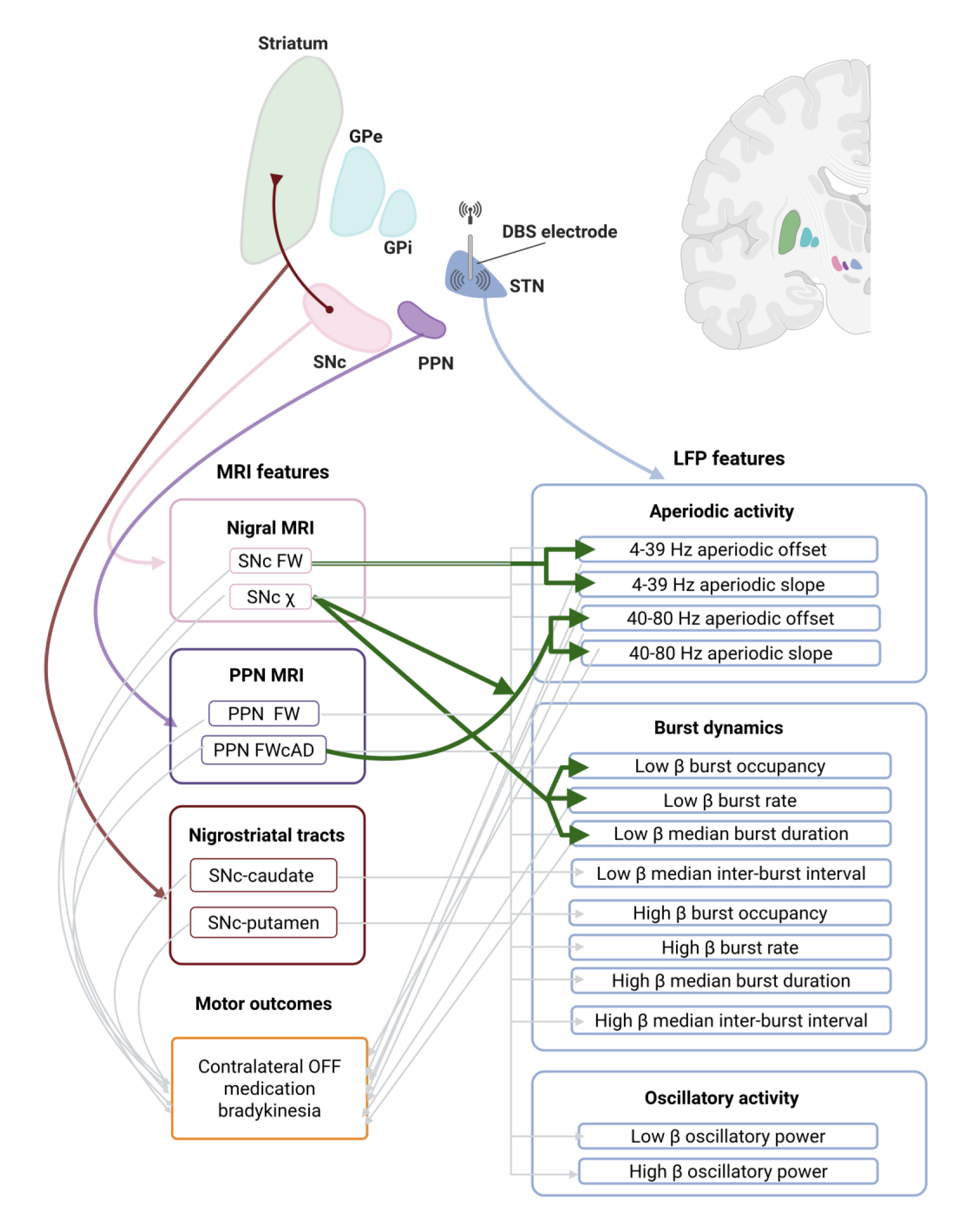
Summary of results. Upper panel provides a schematic overview of the key structures investigated in this study, including the substantia nigra pars compacta (SNc, pink), pedunculopontine nucleus (PPN, purple), subthalamic nucleus (STN, blue), and the nigrostriatal dopaminergic pathway (red). The lower panel summarises all associations tested in the study. Light grey lines indicate associations that were tested but were not statistically significant, whereas solid green lines indicate associations that survived false discovery rate (FDR) correction (q < 0.05). The curved green connection between PPN cAD and the 40–80 Hz aperiodic measures denotes that this relationship was moderated by SNc susceptibility. All models were adjusted for age, residual OFF-medication levodopa exposure, and the relevant global grey matter MRI measure.

## 5. Discussion

Subthalamic LPFs are increasingly used to track and therapeutically respond to the symptomatic state of Parkinson’s disease, but whether they also contain information about the underlying neurodegenerative processes has remained unclear. Here, we show that distinct features of the human STN LFP are differentially associated with in vivo markers of nigral and brainstem tissue pathology. SNc free water was preferentially associated with low-frequency aperiodic activity, SNc susceptibility with low-β burst dynamics, and PPN cAD with high-frequency aperiodic activity. Moreover, the association between PPN cAD and STN aperiodic activity became stronger with increasing SNc susceptibility, consistent with preclinical evidence that PPN influences on the STN depend on dopaminergic integrity^36–38^. These findings extend the biological interpretation of the STN LFP beyond a readout of current motor state, suggesting that its oscillatory and aperiodic components carry partly separable information about the pathological systems contributing to parkinsonian circuit dysfunction. More broadly, they raise the possibility that chronically implanted DBS systems could provide electrophysiological markers of underlying disease biology as well as signals for symptom-responsive therapy.

### Nigral pathology maps onto distinct LFP components

Our findings provide human evidence that variation in nigral tissue pathology is expressed in distinct components of STN electrophysiology. Preclinical models have shown that dopaminergic degeneration alters STN firing, β synchronisation and burst dynamics, but translation of these observations to humans has been limited by the scarcity of datasets combining quantitative markers of nigral pathology with invasive STN recordings. Here, nigral pathology was not expressed uniformly in the STN LFP: SNc susceptibility was preferentially associated with the temporal organisation of low-β bursting, whereas SNc free water mapped onto low-frequency aperiodic activity. This double dissociation extends multimodal MRI evidence that susceptibility and free water capture partly independent aspects of parkinsonian nigral pathology^41^, by showing that their non-redundancy is reflected in downstream human basal ganglia physiology.

The association between SNc susceptibility and low-β burst dynamics may connect iron-sensitive nigral tissue pathology with network abnormalities commonly attributed to dopamine depletion. Longitudinal preclinical studies show that nigrostriatal degeneration is accompanied by increasing STN β activity and prolonged β bursting ^11,13^, linking the temporal organisation of β activity to the progression of dopaminergic pathology. In our human data, susceptibility, which is strongly influenced by nigral iron content, but also by tissue composition and microstructural organisation ^42,43^, was associated with greater low-β burst occupancy, longer bursts and a higher burst rate. These findings suggest that susceptibility-sensitive nigral pathology is expressed downstream in the electrophysiological domain most consistently linked to dopamine depletion: the abnormal temporal organisation of STN β synchronisation.

The association between SNc free water and low-frequency aperiodic activity may identify an additional, non-oscillatory dimension of nigral pathology. Free water is commonly interpreted as reflecting extracellular expansion and microstructural tissue disruption arising from overlapping processes that may include neuronal and axonal loss, neuroinflammatory change and degradation of tissue architecture ^25–27^. It may therefore capture the cumulative structural consequences of nigral degeneration. Such disruption could alter nigrostriatal signalling diffusely, changing the aggregate inputs that shape downstream STN population activity. Influential early models of parkinsonian basal ganglia dysfunction emphasised increased firing rates, altered population activity and STN hyperactivity following dopamine depletion, whereas subsequent human electrophysiological research became increasingly centred on β oscillations^20–22^. Aperiodic decomposition may provide a means of recovering this broader dimension of circuit dysfunction from the human STN LFP. Our findings suggest that the diffuse or cumulative nigral tissue disruption indexed by SNc free water may be preferentially expressed through this broader population-level dimension of parkinsonian circuit dysfunction.

Together, the findings indicate that no single electrophysiological feature provides a complete readout of nigral pathology. Instead, oscillatory burst dynamics and aperiodic population activity appear to reveal complementary aspects of the downstream physiological consequences of nigral tissue change.

#### Impact of PPN degeneration on STN activity

The PPN findings show that the pathological information carried by the STN LFP extends beyond the nigrostriatal system. PPN microstructure mapped selectively onto high-frequency aperiodic activity, indicating that the state of an extranigral brainstem input is expressed in a distinct component of STN population physiology. To our knowledge, this is the first demonstration that in vivo PPN tissue properties relate to human STN electrophysiology. The STN LFP may therefore be understood not as a unitary readout of dopamine loss, but as an integrated physiological signal in which degeneration across convergent neural systems leaves separable signatures.

Higher PPN cAD may reflect loss or reorganisation of PPN integrity, including selective degeneration of grey-matter elements causing partial-volume shifts towards relatively white-matter-dominated voxels, altered organisation of surviving fibre populations, or loss of specific crossing or branching projection systems. We found that higher PPN cAD was associated with a steeper high-frequency aperiodic slope, commonly interpreted as being consistent with a shift in excitation–inhibition balance towards relatively greater inhibition, reduced excitation or longer synaptic timescales^24^. Our findings therefore suggest that altered PPN microstructure may shift the STN towards a less excited population state through several possible routes. Loss of direct cholinergic or glutamatergic PPN–STN projections could reduce excitatory or modulatory drive to the STN^32,33^. The PPN may also influence STN activity indirectly through its excitatory and modulatory inputs to SNc dopaminergic neurons.

Importantly, the interaction with SNc susceptibility supports the broader principle that PPN influences on STN physiology are state-dependent. The association between PPN cAD and high-frequency aperiodic activity was strongest in hemispheres with higher SNc susceptibility, indicating that the electrophysiological expression of PPN tissue alteration changes with the severity of concurrent nigral pathology. This aligns with preclinical evidence that PPN disruption has opposing effects on STN activity depending on dopaminergic integrity: PPN lesions can increase STN firing in dopamine-intact animals, but reduce pathological STN hyperactivity and β activity after nigrostriatal dopamine depletion ^37,38^. One possible explanation is that the relative contribution of PPN pathways changes as dopamine loss advances: effects mediated through surviving SNc neurons become progressively constrained, while direct and other non-dopaminergic PPN influences assume greater importance. The present findings therefore support a network model in which PPN structural alteration does not have a fixed electrophysiological consequence, but is expressed according to the dopaminergic state of the basal-ganglia circuit.

Together, these results suggest that the PPN contributes to STN electrophysiology through a high-frequency aperiodic signature. More broadly, they raise the possibility that extranigral brainstem integrity influences the population state of the STN by modulating the balance of circuit inputs that determine STN excitability. This expands the relevance of the PPN beyond its currently emphasised role in postural instability, gait and freezing ^44–46^. Rather than acting solely as an axial-motor or locomotor node, the PPN may have a broader and currently underappreciated influence over the parkinsonian network state, shaping how nigral pathology is expressed in basal-ganglia physiology. This possibility warrants targeted development of MRI methods capable of characterising the PPN and its projection systems in vivo, particularly approaches that can distinguish free-water change, tissue loss, fibre organisation and partial-volume effects within this small and heterogeneous brainstem region.

#### Impact of LFP activity on symptoms

The clinical analyses suggest that the MRI-linked electrophysiological signatures identified in the preceding sections do not have equivalent relationships with contralateral bradykinesia. The clearest convergence was observed for the low-frequency aperiodic signature linked to SNc free water. Specifically, higher 4–39 Hz aperiodic offset was associated with worse contralateral OFF-medication bradykinesia, and remained the only significant electrophysiological predictor in the exploratory multiparametric model including representative low-frequency aperiodic, low β spectral mass, β-burst and high-frequency aperiodic features. This supports the idea that low-frequency aperiodic activity captures a clinically relevant component of STN dysfunction that is not reducible to conventional β activity.

This interpretation is consistent with recent multicentre evidence that clinical relationships between STN electrophysiology and motor severity extend beyond β rhythms. In a large cross-sectional STN-LFP study, Gerster et al. ^23^ showed that previously inconsistent β-related findings were likely explained in part by limited statistical power and methodological variability. In that study, although low-β activity was positively associated with motor impairment, the effect was modest and required more than 100 patients for stable replication. By contrast, aperiodic offset showed one of the strongest associations with motor severity, and a model combining offset, low beta and low gamma explained more variance than low β alone.

The stronger clinical association with 4–39 Hz aperiodic offset in the current paper is therefore notable. It suggests that the low-frequency aperiodic component provided the most sensitive summary of the STN activity state related to contralateral bradykinesia. This aligns with the broader conclusion of Gerster et al.^23^ that aperiodic activity may provide clinically meaningful information beyond oscillatory β power. In the present data, this aperiodic feature was also the component most clearly linked to SNc free water, suggesting a possible pathway by which nigral tissue disruption is expressed as altered STN population dynamics and, in turn, lateralised motor impairment.

The absence of any relationship between bradykinesia for the PPN cAD-linked high-frequency aperiodic signature is also informative, but should be interpreted differently. Unlike the low-frequency aperiodic signature, which related to contralateral bradykinesia, the PPN-linked high-frequency aperiodic component may reflect a broader STN population state that is not well captured by lateralised limb bradykinesia alone. Given the known involvement of the PPN in axial, gait, postural and arousal-related functions, its electrophysiological consequences may be more apparent in clinical domains not presented here, such as gait variability, freezing, falls, balance, sleep-wake regulation or cognitive-motor control. We are addressing these questions in ongoing analyses using quantitative assessment of these symptom domains.

### Limitations

Our cohort is large for an invasive STN LFP study, but we acknowledge the sample size could raise concerns that the reported findings were driven by individual participants. To address this, we performed extensive sensitivity analyses, all of which supported the robustness of the main results. In addition, while diffusion and susceptibility MRI provide valuable in vivo markers of neurodegeneration, they remain indirect measures of the underlying pathology. The precise biological interpretation of free water, iron accumulation and particularly PPN axial diffusivity remains incompletely understood.

### Conclusions

This study provides the first evidence that distinct features of the human STN LFP are differentially associated with in vivo MRI markers of nigral and brainstem pathology in Parkinson’s disease. SNc free water mapped onto low-frequency aperiodic activity, SNc susceptibility onto low-β burst organisation, and PPN cAD onto high-frequency aperiodic dynamics, with the PPN–STN relationship further shaped by the state of the nigral system. These findings suggest that the STN LFP is not a unitary readout of dopamine loss or motor severity, but an integrated physiological signal in which pathology across interconnected systems is expressed through separable electrophysiological components.

This distinction has important implications for biomarker development. Adaptive DBS has largely focused on physiomarkers that track moment-to-moment symptom severity and can be used to control stimulation, with β activity remaining the dominant candidate. Yet β-based measures explain only part of the variability in motor impairment and are unlikely to capture the full biological heterogeneity of parkinsonian circuit dysfunction^47^. With longitudinal validation, our findings indicate that chronically recorded LFPs may provide not only control signals for symptom-responsive therapy, but also physiological readouts of the distinct neurodegenerative processes shaping basal ganglia dysfunction.

## 6. Methods

### 6.1. Participants

45 participants were recruited, with 33 datasets available for the current analysis. Included participants were six females and 27 males; Mean age=62.11, sd=7.5, who were consecutively recruited from DBS services at Salford Royal Hospital in Manchester and The Walton Centre in Liverpool soon after being approved to proceed for DBS surgery.

### 6.2 Data Collection

#### 6.2.1 Pre-surgery

All participants underwent multi-modal MRI scanning on a Philips 5.7.1.6 prior to surgery. Images were collected while participants were ON medication.

##### 6.2.1.1 MRI Acquisition Parameters

Four T1-weighted images (for subsequent averaging) were obtained using an MP-RAGE sequence with a repetition time of 2.3 milliseconds, an echo time of 11.39 milliseconds, and a flip angle of 8°. The SENSE factor was 2.14, the field of view was 160.3 x 268.8 x 268.8, and the voxel size was 0.7 × 0.7 × 0.7 mm³.

Diffusion-weighted images were acquired using a 2D diffusion-weighted, spin-echo, echo planar imaging sequence with a repetition time of 5.6 milliseconds, echo time of 80 milliseconds, and a flip angle of 90°. The voxel size was 2.5 x 2.5 x 2.5, slice thickness was 2.5 mm, and the field of view was 240 x 240 x 150 mm. Diffusion weighting was applied in 112 directions: 8 directions with a b-value of 300 s/mm², 32 directions with a b-value of 1000 s/mm², and 64 directions with a b-value of 2000 s/mm². Additionally, nine B0 images were acquired without diffusion weighting. A reverse phase-encoded image with no diffusion weighting was also collected for use in fsl’s topup.

Multi-echo GRE images were acquired using a sequence with 3 echoes at echo times of 5.4, 14.1, and 22.8 milliseconds. The flip angle was 17°, and the voxel size was 0.68 × 0.68 × 1 mm³. The field of view and matrix size were 228.48 x 228.48 x 140 and 336 x 336 x 140 respectively.

##### 6.2.1.2 Image preprocessing

T1-weighted imaging: The four 0.7mm T1w images were affine registered to the first scan and averaged together to improve the signal-to-noise ratio. T1-weighted images were bias-field corrected and intensity-scaled to a range of 0–255. Skull stripping was performed using the SynthStrip Toolbox.

Diffusion-weighted imaging: Data were first denoised (dwidenoise, MRtrix3) and then corrected for eddy-current induced distortions and motion using FSL’s Topup and Eddy toolboxes48,49. The resulting images were then bias field corrected (dwibiascorrect ants) and upsampled to an isotropic resolution of 1 mm (MRTrix3).

Tractography: Upsampled diffusion data were used to generate response functions for white matter, grey matter, and cerebrospinal fluid, using the dhollander algorithm implemented in MRtrix3 50.

Multi-shell, multi-tissue constrained spherical deconvolution (MSMT-CSD) was performed using the group averaged tissue response functions to compute fibre orientation distributions (FODs). Multi-tissue intensity normalization was applied across the FOD images to ensure inter-tissue comparability (mtnormalise).

A five-tissue-type segmentation was generated from the T1-weighted anatomical image (5ttgen fsl) and coregistered to the non-diffusion-weighted DWI image, b = 0, using ANTs. The subcortical grey matter compartment was modified (using 5ttedit) to include the bilateral SNc, substantia nigra pars reticulata (SNr), STN, red nucleus, and globus pallidus internus and externus from the HybraPD atlas; the PPN from the human PPN atlas; and the caudate nucleus, putamen, and thalamus from FSL FIRST. This enabled streamlines to enter and propagate within these regions while constraining their subsequent propagation, thereby permitting region-wise connectivity estimation.

Whole brain tractography was performed using tckgen (MRtrix3) with the iFOD2 algorithm, dynamically seeding from the intensity-normalised white matter FOD image to generate a 20 million streamline whole brain tractogram for each individual (backtracking enabled; maximum length = 250 mm). The default minimum streamline length (2 x voxel size) was retained to allow reconstruction of short-range connections between closely adjacent structures. Whole brain tractogram streamline densities were then matched to the underlying diffusion signal by SIFT2-filtering, producing a weight value per streamline51.

Tckedit was used to isolate streamlines connecting the SNc and striatum, along with their corresponding weights, separately for the caudate nucleus and putamen from the whole brain tractograms. To exclude anatomically implausible, spurious, or interhemispheric crossing streamlines, a maximum length threshold was applied, defined as twice the Euclidean distance between the ROI centroids. In addition, a bounding box encompassing all subcortical structures was used to further constrain the tractography results.

Connection strength between regions was quantified as the sum of streamline weights, scaled by the subject specific SIFT proportionality coefficient (mu) to permit quantitative comparisons between individuals52 . This metric has been taken to reflect the capacity of a bundle to transmit information (known as ‘fibre bundle capacity’).

###### Freewater

Free water and free water-corrected diffusion metrics were estimated using a multi-shell bi-tensor diffusion model implemented in NiPy53 . The preprocessed multi-shell diffusion data (as described above) were used as input to a voxelwise two-compartment model in which the diffusion signal is represented as the sum of an anisotropic tissue compartment (modelled with a diffusion tensor) and an isotropic free-water compartment. Both the tissue tensor and the free-water fraction were fitted from the data, allowing the free-water signal fraction to vary across voxels. Voxelwise free water maps (fractional volume of the isotropic compartment) and free water-corrected tissue tensors were obtained, and free water-corrected axial diffusivity (cAD) maps were then derived from the principal eigenvalue of the free water-corrected tensor.

Quantitative susceptibility mapping (QSM): To generate a brain mask, the first echo of the magnitude image was extracted and bias-field corrected. Due to suboptimal performance of standard skull-stripping tools on magnitude images, an affine registration was performed between the unstripped T1 and the magnitude image, and the resulting transform was used to bring the T1-derived brain mask into magnitude space.

Pre-processing was performed using the SEPIA toolbox 54. The T1w brain mask was applied consistently throughout the QSM pipeline. Phase unwrapping was carried out using the 3D Laplacian method implemented in the MEDI algorithm 55. Background field removal was conducted using the PDF method 56, with local field refinement performed using a 3D fourth-order polynomial fit (refine_order = 4), and no additional erosion applied before or after refinement (erode_radius = 0; erode_before_radius = 0). The dipole inversion was completed using the MEDI algorithm57, with cerebrospinal fluid (CSF) used as the reference tissue

Next, a hybrid T1-QSM image was generated. This approach was motivated by evidence that combining T1-weighted and susceptibility contrast yields higher-quality templates than T1-only methods 58, by preserving anatomical context while enhancing contrast in deep grey matter structures that are better defined on QSM. As such, an affine registration was performed using ANTs to align the skull-stripped magnitude images with T1w images. Following Hanspach et al.58, we next created a hybrid T1-QSM image according to the following formula:

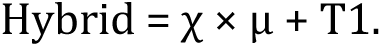

The scaling factor μ was determined as the ratio between the maximum T1 intensity and the maximum susceptibility (χ) value within the basal ganglia (BG) region. This approach is intended to preserve the polarity of paramagnetic and diamagnetic contrasts in the hybrid image. To obtain maximum T1 intensity values, a BG mask was created from each participant’s segmented T1w image (via FSL). T1w and QSM intensity values at the 99.5th percentile were used to define the upper limit for μ. Subsequently, a range of μ values within this limit were tested. The final μ was selected based on visual inspection, prioritizing the value that provided the clearest anatomical delineation of the midbrain.

##### 6.2.1.3 Atlas registration and region-of-interest (ROI) delineation

SNc and PPN were identified using the HybrA-PD atlas59 and human PPN atlas 60 . Whole brain GM masks were generated from fsl’s fast of the processed T1 image. Each individual’s hybrid image was registered to the HybrA-PD and human PPN templates. Inverse transforms were used to map the ROI masks into each individual’s QSM space to extract susceptibility measures. To create the same ROI masks in DWI space, the transforms were concatenated from template-to-QSM, QSM-to-T1w, and finally T1w-to-b0 images.

#### 6.2.2 Post surgery

As soon as possible after participants had recovered from surgery, they were invited to complete the following tasks in the OFF medication and OFF DBS state. Medication withdrawal protocols were individualized and supervised by the treating neurologist to ensure participant safety and to achieve a suitable OFF medication state at the time of data collection. Only OFF medication outcomes are reported here, though the full protocol included an ON medication session.

Medication withdrawal was limited to a maximum of 24 hours prior to participation. As a result, dopaminergic medications with long half-lives (e.g., dopamine agonists) were still present in the system during the OFF medication condition. To account for this, the residual levodopa equivalent dose (LED) was calculated via a dopaminergic pharmacokinetics tool 61. Subsequently, LED at the time of each session was controlled for in all statistical analyses (see statistical analysis section for further detail).

For resting state recordings, participants were seated in a dark room and instructed to remain as still as possible for 8 minutes to minimize movement-related artifacts. LFPs were recorded using the Medtronic Percept BrainSense system.

Participants also underwent evaluation with the MDS-UPDRS Part III. The full protocol also included a gait and balance assessment and computer-based and pen-paper cognitive tasks (these are not reported in the current paper).

##### 6.2.2.1 Signal Processing

LFP signals were recorded at 250Hz, upsampled to 500 Hz and segmented into 4-second epochs with no overlap. For the exploration of the power spectrum, the data were bandpass filtered between 4 and 39 Hz and zero-padded to a total length of 4 seconds. Spectral estimation was performed using the multitaper method with ±2 Hz frequency smoothing applied around each frequency bin.

To isolate the oscillatory component of the power spectrum, the specparam toolbox in python was used to decompose the signal. As a first step both the fixed and the knee models were used to extract the fit from each participant and condition. Visual inspection was used to decide on the best model for each case, as well as the quantitative model fit information (R2 and MAE). Finally the model that best fit the data in each case was used for follow up analyses. The oscillatory component was derived by dividing the original spectrum by the aperiodic component, using the formula

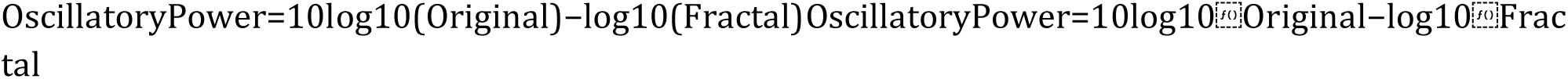

The oscillatory spectrum was bandpass filtered into low and high β (13–20 Hz and 21-30 Hz respectively). Power within each band was extracted and converted to decibel units using a logarithmic transform:

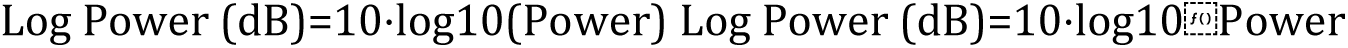

To isolate the aperiodic component on low (4-39Hz) and high frequency bands (40-80Hz) the same approach to the one above was taken, however, in this case the aperiodic component was estimated only using the fixed method since a knee artifact would only be observable in a wider power spectrum. The metrics derived from both low and high frequency band aperiodic fits were the offset and slope.

##### 6.2.2.2 Burst computation

To perform burst computation the original signal which included both the aperiodic and oscillatory component was used.

To extract burst metrics, the ON- and OFF-medication recordings from each participant were first standardised together by dividing both recordings by the standard deviation of the concatenated ON and OFF signals. The two recordings were then processed independently.

For each recording, the signal was bandpass filtered within the frequency band of interest using a two-pass finite impulse response (FIR) filter. The amplitude envelope was obtained by applying the Hilbert transform to the filtered signal. The envelope was subsequently smoothed using a Gaussian window, and slow baseline fluctuations were removed by subtracting a long-timescale Gaussian-smoothed baseline. The detrended envelope was then converted to a robust z-score according to:

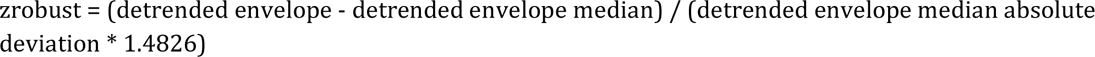

Bursts were defined as periods during which the robust z-score exceeded a threshold of 2. Bursts shorter than two oscillatory cycles of the centre frequency were excluded from further analysis. Once the burst were detected the following metrics were computed: Burst occupancy (percentage of time spent in bursts), burst rate (bursts/s), median burst duration, median inter-burst interval (time between consecutive bursts), and median burst magnitude above threshold.

### 6.3 Statistical analysis

Statistical analyses were performed at the hemisphere level to test whether OFF-medication STN electrophysiological features were associated with quantitative MRI markers of nigral and brainstem pathology and with contralateral OFF-medication bradykinesia. Electrophysiological measures comprised aperiodic offset and slope estimated separately over lower (4–39 Hz) and higher (40–80 Hz) frequency ranges, aperiodic-adjusted low-β and high-β oscillatory spectral mass, and low-β and high-β burst-temporal measures. Burst measures included occupancy, rate, median duration and median inter-burst interval. Imaging predictors comprised ipsilateral SNc free water, SNc susceptibility, PPN free-water-corrected axial diffusivity (cAD), PPN free water, and quantitative measures of the ipsilateral SNc–caudate and SNc– putamen pathways.

Linear mixed-effects models were fitted to hemisphere-level observations, with participant included as a random intercept to account for the non-independence of left- and right-hemisphere measurements. Continuous predictors, outcomes and covariates were standardised within the complete-case sample used for each model; reported β coefficients therefore represent standardised effect estimates. Unless otherwise specified, models were adjusted for age and estimated residual OFF-medication levodopa exposure. Residual levodopa exposure was estimated from the dose, timing and pharmacokinetic properties of dopaminergic medication remaining after the medication-withdrawal period.

#### Nigral MRI–electrophysiology analyses

Associations between SNc tissue properties and ipsilateral STN electrophysiology were examined in separate models for SNc free water and SNc susceptibility. SNc free-water models were additionally adjusted for whole-brain grey-matter free water, whereas SNc susceptibility models were adjusted for whole-brain grey-matter susceptibility. Ipsilateral SNc-caudate and SNc–putamen tract measures were examined in separate pathway-level models without a matched whole-brain MRI covariate.

Electrophysiological outcomes were grouped into prespecified families: lower-frequency aperiodic parameters, higher-frequency aperiodic parameters, low-β oscillatory spectral mass, high-β oscillatory spectral mass, low-β burst-temporal measures and high-β burst-temporal measures. Each MRI predictor was tested separately against the outcomes within these families.

SNc voxel-retention quality control was applied to analyses in which SNc free water was included either as the primary predictor or as a covariate. The primary threshold required at least 10% of SNc voxels to be retained in the relevant hemisphere. Sensitivity analyses repeated the principal SNc free-water models using a stricter 30% voxel-retention threshold.

#### PPN MRI–electrophysiology analyses

PPN cAD and PPN free water were examined separately as predictors of the same STN electrophysiological outcomes. Primary PPN cAD models were adjusted for whole-brain grey-matter cAD, and primary PPN free-water models were adjusted for whole-brain grey-matter free water. To determine whether PPN–STN associations were independent of concurrent nigral tissue variation, secondary models separately added ipsilateral SNc susceptibility or SNc free water. A modality-matched model additionally adjusted the PPN cAD association for ipsilateral SNc cAD.

#### PPN × nigral-pathology interaction analyses

Based on the observed association between PPN cAD and 40–80 Hz aperiodic activity and a prespecified directional hypothesis derived from preclinical evidence, follow-up models tested whether ipsilateral SNc susceptibility moderated the association between PPN cAD and 40–80 Hz aperiodic offset and slope. These models included the PPN cAD main effect, the SNc susceptibility main effect, their interaction, age, estimated residual OFF-medication levodopa exposure, the matched whole-brain grey-matter cAD measure and a participant random intercept.

Directional one-tailed tests were used for the prespecified positive PPN cAD × SNc susceptibility interaction. Specificity was examined using corresponding models in which SNc free water or SNc cAD replaced SNc susceptibility as the nigral moderator, and models in which PPN free water replaced PPN cAD as the PPN predictor. SNc susceptibility was treated as a continuous moderator in all inferential models; grouped values shown in interaction plots were used for visualisation only.

#### Electrophysiology–bradykinesia analyses

To test the clinical relevance of the MRI-linked electrophysiological signatures, mixed-effects models examined associations between ipsilateral STN electrophysiological features and contralateral OFF-medication bradykinesia. Eight features were carried forward: 4–39 Hz aperiodic offset and slope; low-β burst occupancy, median duration and rate; 40–80 Hz aperiodic offset and slope; and low-β oscillatory spectral mass. Models were adjusted for age and estimated residual OFF-medication levodopa exposure and included participant as a random intercept. No MRI covariate or SNc voxel-retention filter was applied to these LFP-only clinical models.

The seven features identified from the preceding MRI–electrophysiology analyses were treated as the primary carried-forward LFP family. Benjamini–Hochberg corrections were calculated both across these seven tests and, as a secondary analysis, within each MRI-linked electrophysiological signature. Low-β oscillatory spectral mass was examined as a targeted benchmark analysis because of previous evidence relating periodic low-β activity to motor impairment; it was not added retrospectively to the original seven-test MRI-linked FDR family.

An exploratory multiparametric mixed-effects model was used to assess whether representative electrophysiological features contributed independent explanatory value for contralateral bradykinesia. Continuous terms were standardised within the common complete-case sample, and variance inflation factors were calculated to assess collinearity. Benjamini–Hochberg correction was applied across the electrophysiological terms included in this model.

#### Direct MRI–bradykinesia analyses

Separate mixed-effects models tested whether ipsilateral SNc free water, SNc susceptibility, SNc cAD, PPN cAD, PPN free water, SNc–caudate pathway strength or SNc–putamen pathway strength was associated with contralateral OFF-medication bradykinesia. Models were adjusted for age, estimated residual OFF-medication levodopa exposure and, for regional MRI predictors, the corresponding whole-brain grey-matter MRI measure. SNc voxel-retention quality control was applied to models containing free-water or cAD measures according to the procedure described above.

#### Multiple-comparison correction and sensitivity analyses

False discovery rate was controlled using the Benjamini–Hochberg procedure. For MRI–electrophysiology main-effect analyses, correction was applied within each MRI predictor and prespecified electrophysiological outcome family. For PPN analyses, correction was performed separately within each PPN predictor, covariate specification and electrophysiological family. The two targeted PPN cAD × SNc susceptibility interaction tests were evaluated as a prespecified directional family. The primary carried-forward electrophysiology–bradykinesia analyses were corrected across the seven MRI-linked LFP features, with additional within-signature corrected values reported as sensitivity analyses. Direct MRI–bradykinesia models were corrected separately across the tested MRI predictors.

Model stability was evaluated using several sensitivity analyses. Models were inspected for singular fits, and fixed-effects models excluding the participant random intercept were fitted as sensitivity analyses. Variance inflation factors were calculated to assess multicollinearity. Leave-one-hemisphere-out analyses sequentially removed each hemisphere, re-standardised all model variables within the reduced sample and refitted the model to assess coefficient stability, sign reversals and dependence on individual observations. Matched-sample analyses were performed where relevant to determine whether differences between MRI predictors or covariate specifications were attributable to variation in complete-case samples.

