## Supplemental for "Distinct nigral and brainstem pathology markers map onto separable subthalamic electrophysiological signatures in Parkinson’s disease"

### Supplemental Material

Supplemental Table 1

| Measure family | Measure | Mean ± SD |
|---|---|---|
| **Nigral MRI** | | |
| N=31, H=60 | SNc free water | 0.403 ± 0.091 |
| N=26, H=51 | SNc susceptibility | 0.164 ± 0.051 |
| **PPN MRI** | | |
| N=31, H=61 | PPN free water | 0.294 ± 0.072 |
| | PPN FWc axial diffusivity | 8.421 ± 0.599 |
| **Nigrostriatal tracts** | | |
| N=33, H=65 | SNc–caudate tract SW μ | 0.258 ± 0.140 |
| | SNc–putamen tract SW μ | 0.343 ± 0.181 |
| **Aperiodic activity** | | |
| N=32, H=63 | 4–39 Hz aperiodic offset | 0.592 ± 0.676 |
| | 4–39 Hz aperiodic slope | 1.273 ± 0.387 |
| | 40–80 Hz aperiodic offset | 2.795 ± 1.412 |
| | 40–80 Hz aperiodic slope | 2.730 ± 0.715 |
| **Aperiodic-corrected oscillatory activity** | | |
| N=31, H=61 | Low-β oscillatory log SM | 12.552 ± 3.71 |
| | High-β oscillatory log SM | 15.118 ± 3.390 |
| **β-burst dynamics** | | |
| N=33, H=65 | Low-β burst occupancy | 0.051 ± 0.014 |
| | Low-β burst rate | 0.223 ± 0.052 |
| | Low-β median burst duration | 0.200 ± 0.018 |
| | Low-β median inter-burst interval | 3.184 ± 1.649 |
| | High-β burst occupancy | 0.062 ± 0.015 |
| | High-β burst rate | 0.378 ± 0.099 |
| | High-β median burst duration | 0.145 ± 0.014 |
| | High-β median inter-burst interval | 1.690 ± 0.537 |

Table 1. Values are reported as mean ± standard deviation (SD). SNc, substantia nigra pars compacta; PPN, pedunculopontine nucleus; STN, subthalamic nucleus; FWcAD, free-water-corrected axial diffusivity; SW μ, tract-level μ derived from spherical mean diffusion modelling; β, beta frequency band. Participant and hemisphere counts vary across measures due to differences in data availability and quality-control exclusions.

### Section 1: Sensitivity analyses for SNc MRI predictors of STN electrophysiological features

We performed additional sensitivity analyses to assess the robustness of the Section 4.1 models relating SNc MRI markers to OFF-medication STN electrophysiological features. These analyses focused on the five effects that survived FDR correction in the primary mixed-effects models: three associations between SNc susceptibility and low-beta burst dynamics, and two associations between SNc free water and 4–39 Hz aperiodic activity.

Leave-one-hemisphere-out analyses showed strong stability of the FDR-significant Section 6.1 effects. The direction of all effects was preserved when individual hemispheres were removed in turn, with no sign reversals observed. Four of the five effects remained nominally significant in every leave-one-hemisphere refit, including both SNc free-water associations with 4–39 Hz aperiodic activity and the two strongest SNc susceptibility associations with low-beta burst dynamics. The remaining association, between SNc susceptibility and low-beta burst rate, remained nominally significant in 50 of 51 refits (Supp Table 1).

| MRI metric | LFP metric | Primary β | Primary p | Primary q | Leave-one-hemisphere β range | LOO p < .05 | Matched β | Matched p |
|---|---|---|---|---|---|---|---|---|
| SNc FW | 4–39 Hz aperiodic slope | 0.355 | .00984 | .0384 | 0.296 to 0.412 | 60/60 | 0.391 | .00735 |
| SNc FW | 4–39 Hz aperiodic offset | 0.258 | .0192 | .0384 | 0.223 to 0.314 | 60/60 | 0.221 | .0533 |
| SNc χ | Low-beta % burst time | 0.591 | $5.87 \times 10^{-6}$ | $4.69 \times 10^{-5}$ | 0.527 to 0.647 | 51/51 | 0.578 | $1.09 \times 10^{-5}$ |
| SNc χ | Low-beta median burst duration | 0.508 | .000611 | .00244 | 0.416 to 0.567 | 51/51 | 0.494 | .000994 |

| | | | | | | | | |
|---|---|---|---|---|---|---|---|---|
| SNc χ | Low-beta burst rate | 0.331 | .0135 | .0360 | 0.251 to 0.435 | 50/51 | 0.310 | .0212 |

Second, we repeated the SNc free-water models using a stricter SNc voxel-retention QC threshold of 30%. Relative to the primary 10% threshold, the 30% threshold reduced the complete-case sample for the aperiodic and burst-temporal models from 60 to 54 hemispheres. The primary SNc free-water associations with low-frequency aperiodic activity were retained. At the 30% threshold, higher SNc free water remained associated with steeper 4–39 Hz aperiodic slope (β = 0.389, t = 2.92, p = .0053) and greater 4–39 Hz aperiodic offset (β = 0.263, t = 2.58, p = .0132).

Third, we performed matched-sample analyses to test whether the apparent dissociation between SNc free-water and susceptibility effects could be explained by differences in sample composition. For each electrophysiological outcome, SNc free-water and susceptibility models were repeated in the same complete-case hemisphere sample, requiring available SNc free water, SNc susceptibility, both matched-modality global covariates, and passage of the primary 10% SNc voxel-retention threshold. In this matched sample, SNc free water remained associated with 4–39 Hz aperiodic slope (β = 0.391, t = 2.81, p = .00735, N = 50 hemispheres). The association with 4–39 Hz aperiodic offset remained positive but marginally missed conventional significance (β = 0.221, t = 1.99, p = .0533, N = 50 hemispheres). SNc susceptibility remained strongly associated with low-beta percentage burst time (β = 0.578, t = 4.98, p = $1.09 \times 10^{-5}$, N = 50 hemispheres), low-beta median burst duration (β = 0.494, t = 3.53, p = .000994, N = 50 hemispheres), and low-beta burst rate (β = 0.310, t = 2.39, p = .0212, N = 50 hemispheres). Thus, the matched-sample analyses supported the primary dissociation between SNc free-water and susceptibility effects (Supp Table 1).

Finally, across the primary models, no nominally significant mixed-effects models were singular, and variance inflation factors did not indicate problematic collinearity. Together, these sensitivity analyses support the robustness of the Section 6.1 findings.

### Supplementary Results: Section 2

We performed additional sensitivity analyses to assess the robustness of the Section 4.2 models relating PPN MRI markers to OFF-medication STN electrophysiological features.

These analyses focused on the core 40–80 Hz aperiodic effects: the positive associations between PPN FWcAD and 40–80 Hz aperiodic offset and slope, their stability after adjustment for ipsilateral SNc MRI markers, the opposing associations with PPN free water, and the PPN FWcAD × SNc susceptibility interaction.

Leave-one-hemisphere-out analyses showed strong stability of the PPN FWcAD findings. Across whole-brain-adjusted, SNc susceptibility-adjusted, and SNc free-water-adjusted models, the positive direction of the PPN FWcAD effect was preserved when individual hemispheres were removed in turn, and all refits remained significant across all covariate specifications. These results indicate that the core PPN FWcAD effects were not driven by a single hemisphere.

| PPN metric | LFP metric | Whole-brain β | Whole-brain p | SNc-adjusted robustness | Leave-one-hemisphere β range | LOO p < .05 | VIF range |
|---|---|---|---|---|---|---|---|
| PPN FWcAD | 40–80 Hz aperiodic offset | 0.347 | .00614 | Retained after SNc χ and SNc FW adjustment; βs = 0.330–0.362, ps = .00841–.0106, qs = .0169–.0213 | 0.277 to 0.395 | 172/172 | 1.07–1.10 |
| PPN FWcAD | 40–80 Hz aperiodic slope | 0.349 | .00576 | Retained after SNc χ and SNc FW adjustment; βs = 0.333–0.360, ps = .00846–.00957, qs = .0169–.0213 | 0.293 to 0.390 | 172/172 | 1.07–1.10 |
| PPN FW | 40–80 Hz aperiodic offset | −0.264 | .0384 | Strengthened after SNc χ adjustment but not retained after SNc FW adjustment; SNc χ: β = −0.386, p = .00734, q = .0214; SNc FW: β = −0.221, p = .149, q = .343 | −0.432 to −0.222 | 114/122 | 1.17–1.90 |

| | | | | | | | |
|---|---|---|---|---|---|---|---|
| PPN FW | 40–80 Hz aperiodic slope | −0.265 | .0378 | Strengthened after SNc χ adjustment but not retained after SNc FW adjustment; SNc χ: β = −0.367, p = .0107, q = .0214; SNc FW: β = −0.208, p = .172, q = .343 | −0.409 to −0.225 | 115/122 | 1.17–1.90 |

The PPN FWcAD × SNc susceptibility interaction was positive for both 40–80 Hz aperiodic offset in the primary analysis. The planned one-tailed interaction remained below $p < .05$ in 49 of 51 leave-one-hemisphere refits for both 40–80 Hz aperiodic offset and slope. These results support the interpretation that the PPN FWcAD–STN aperiodic relationship was strongest in hemispheres with higher SNc susceptibility (Supp Table 3).

| **Interaction** | **LFP metric** | **β** | **Two-tailed p** | **One-tailed p** | **Leave-one-hemisphere β range** | **One-tailed LOO p < .05** | **VIF** |
|---|---|---|---|---|---|---|---|
| PPN FWcAD × SNc χ | 40–80 Hz aperiodic offset | 0.311 | .0578 | .0289 | 0.260 to 0.392 | 49/51 | 1.60 |
| PPN FWcAD × SNc χ | 40–80 Hz aperiodic slope | 0.313 | .0546 | .0273 | 0.266 to 0.376 | 49/51 | 1.60 |

Across the Section 4.2 sensitivity analyses, variance inflation factors did not indicate problematic collinearity, with no models exceeding VIF > 5. Several significant 40–80 Hz mixed-effects models were singular, consistent with negligible participant-level random-intercept variance; however, equivalent fixed-effects models yielded comparable or identical estimates. Together, these sensitivity analyses support the robustness of the Section 4.2 findings.
